# Semantic and population analysis of the genetic targets related to COVID-19 and its association with genes and diseases

**DOI:** 10.1101/2022.09.16.508278

**Authors:** Louis Papageorgiou, Eleni Papakonstantinou, Io Diakou, Katerina Pierouli, Konstantina Dragoumani, Flora Bacopoulou, George P Chrousos, Elias Eliopoulos, Dimitrios Vlachakis

## Abstract

SARS-CoV-2 is a coronavirus responsible for one of the most serious, modern worldwide pandemics, with lasting and multi-faceted effects. By late 2021, SARS-CoV-2 has infected more than 180 million people and has killed more than 3 million. The virus gains entrance to human cells through binding to ACE2 via its surface spike protein and causes a complex disease of the respiratory system, termed COVID-19. Vaccination efforts are being made to hinder the viral spread and therapeutics are currently under development. Towards this goal, scientific attention is shifting towards variants and SNPs that affect factors of the disease such as susceptibility and severity. This genomic grammar, tightly related to the dark part of our genome, can be explored through the use of modern methods such as natural language processing. We present a semantic analysis of SARS-CoV-2 related publications, which yielded a repertoire of SNPs, genes and disease ontologies. Population data from the 100Genomes Project were subsequently integrated into the pipeline. Data mining approaches of this scale have the potential to elucidate the complex interaction between COVID-19 pathogenesis and host genetic variation; the resulting knowledge can facilitate the management of high-risk groups and aid the efforts towards precision medicine.

## Introduction

*Coronaviridae* is a family of enveloped viruses with a single-stranded RNA genome roughly 25 to 32 kb long. The size of the virion ranges from 118 to 136 nm, while its surface is studded with the characteristic, large spike (S) glycoprotein. The viral family is further divided into subfamilies *Orthocoronavirinae* and *Letovirinae*, with the latter comprising of a single genus, *Alphaletovirus*. The *Orthocoronavirinae* subfamily circumscribes four non-monotypic genera, namely *Alpha*-, *Beta*-, *Gamma*- and *Deltacoronavirus* (1). Alpha- and betacoronaviruses commonly infect mammals, gammacoronaviruses mainly infect avian species, while deltacoronaviruses infect mammals and birds (2). The effects of human coronavirus infection can range from mild, such as in the case of HCoV-229E, to potentially life-threatening, such as in the case of the Middle East respiratory syndrome coronavirus (MERS-CoV) and severe acute respiratory syndrome coronavirus (SARS-CoV-1 and 2) (3).

The standard coronavirus virion comprises of the membrane, envelope and spike proteins, which are all embedded in the viral envelope, as well as the nucleocapsid protein, which interacts with the viral RNA at the virion’s core (4). The large coronavirus genome possesses untranslated regions at both ends; two large ORFS at the 5’ end, ORF1a and ORF1b, code for non-structural proteins necessary for the formation of the replication and transcription complex (RTC), while ORFs encoding structural and accessory proteins are transcribed from the 3’ end (5). During infection, the coronavirus spike (S) protein mediates binding to specific cellular receptors. For example, both the betacoronaviruses SARS-CoV and SARS-CoV-2 recognize the angiotensin-converting enzyme 2 (ACE2) (6), while other betacoronaviruses like MERS-CoV and HKU4 recognize the dipeptidyl peptidase 4 (DPP4) (7). The S protein is a homotrimeric, class I fusion glycoprotein, forming petal-shaped projections on the virion’s surface (8). In some coronaviruses, S is cleaved during the maturation process while in others, including SARS-CoV, S is arranged into two domains, S1 and S2, with different functions (9, 10). Within the surface-exposed S1 domain lies the receptor-binding domain (RBD) responsible for interaction with the host cell receptor while the transmembrane S2 domain mediates fusion between viral and host cell membranes (11). In a study by Wang *et al*., the SARS-CoV spike protein, through interaction with murine macrophages, was found to induce IL-6 cytokines and release of TNF-α (12). Interleukin-6 (IL-6) plays a key role in the innate and acquired immune response, inducing the acute-phase response after the occurrence of infection and inflammation (13). In a preprint, Hsu *et al*. proposed that the SARS-CoV-2 spike protein induces significant NF-κB activations as well as production of pro-inflammatory cytokines (14). The described mechanism of action is the stimulation of the MAPK-NF-κB axis through the binding of the S protein to the ACE2 receptor, resulting in the release of cytokines (14). Furthermore, in a recent preprint, modeling and docking studies highlighted a potential interaction between the SARS-CoV-2 spike glycoprotein and nicotinic acetylcholine receptors (nAChRs), proposing an underlying mechanism participating in severe COVID-19 (15).

Since being declared in March 2020, the ongoing COVID-19 pandemic has affected countries on a near global scale, with more than 248 million confirmed cases and more than 5 million deaths by November 2021 (https://www.who.int/), challenging healthcare systems, economies and communities in multiple ways. COVID-19 exhibits strong heterogeneity when it comes to clinical representation, ranging from asymptomatic to severe disease affecting multiple organs (16). Influenza-like symptoms tend to be prevalent, as the main sites of infection are the upper and lower respiratory tract, however other organs such as the heart of kidneys can be affected as sites of ACE2 expression (17). Factors which impact the risk and severity of COVID-19 are continuously being investigated. A 2021 meta-analysis of more than 17 million patient data highlighted common variables linked to adverse outcome, such as older age, severe obesity and active cancer (18, 19). One drug, remdesivir, has been approved by the FDA for treatment of COVID-19 while investigational therapies, such as monoclonal antibodies, are being explored. Chen *et al*. reported the isolation of two lead IgG1 monoclonal antibodies which effectively blocked the binding between ACE2 and the SARS-CoV-2 RBD (20). Out of a set of neutralizing antibodies isolated by Wu *et al*., two antibodies, B38 and H4, effectively blocked the binding of the RBD to ACE2 (20). CB6, a specific human monoclonal antibody isolated by Shi *et al*., was shown to hinder SARS-CoV-2 infection in vitro as well as in rhesus monkeys, by targeting an epitope that overlaps with ACE2 binding sites in the SARS-CoV-RBD (21). Non-RBD monoclonal antibodies are also investigated (22).

To curtail the damaging effects of COVID-19 and expedite herd immunity, the scientific community raced to develop vaccines against SARS-CoV-2. Currently available vaccines rely on the spike protein as an immunogen because of its key roles during viral entry; the first category, mRNA and adenoviral vector vaccines, provide genetic information for spike protein synthesis, while the second category, inactivated vaccines, constitute protein-based strategies (23). By November 2021, more than 53% of the world population has received at minimum one dose of a COVID-19 vaccine (24). Nevertheless, vaccine hesitancy is a widespread phenomenon, as evidenced by data stemming from behavior analysis conducted by the Imperial College of London (25). In surveys about citizens’ willingness to get vaccinated against COVID-19 in Germany, France, Italy, Australia, Spain and Japan, the share of the surveyed population who were unvaccinated and unwilling to get vaccinated ranged between 12-22% (25). Through global circulation, SARS-CoV-2 variants have and will continue to emerge as a result of selective pressure and continuous viral replication within the population of hosts. One likely selective pressure is for mutations which improve intrinsic fitness, such as in the case of the D164G substitution in the spike protein (26). Increased infection in the upper airway due to D164G has allowed the variant to dominate over the wild-type virus (26, 27). The Delta variant (B.1.617.2), which is becoming the dominant strain globally according to WHO, has been shown to be eightfold less sensitive to vaccine-elicited antibodies in comparison to the wild-type Wuhan-1 bearing D164G in vitro (26). Therefore, the global scientific community is called to keep a close eye on the ever-changing landscape of the SARS-CoV-2 mutational landscape and its potential effects on the vaccines’ effectiveness.

Genome-wide association studies (GWAS) are an important tool in the investigation of disease pathogenesis and enable the characterization of relevant single nucleotide polymorphisms (SNPs) (28). Furthermore, genetic variants which are linked to diseases can shape a polygenic risk score, which characterizes the individual’s susceptibility to certain diseases (29, 30). As it has been evidenced, polymorphisms occurring in regions that do not code for proteins are frequent and can have equally potent effects (31, 32). Therefore, when exploring the variation of the individual’s genetic makeup, it would be unwise to limit ourselves to the coding regions of the human genome. When analyzing genetic variation under the scope of infection and disease risk, “genomic grammar” can be an appropriate term, since it is not limited to the gene but encompasses factors related to the dark part of the genome which have only recently begun to be investigated (32).

The 1000 Genomes Project provides an invaluable pool of whole genome sequencing data, with a goal of constructing an inventory of genetic variations within the human genome (33). Genomes of more than 2.500 individuals have been mapped for genetic variation (34). During the project’s analysis, a specific allele frequency is assigned to each pinpointed variant, calculated by dividing the number of the allele’s occurrence in the population by the total sum of copies of all the alleles at the genetic locus of interest. Data provided by the 1000 Genomes Project include – among others - the general allele frequency of the determined variants and the corresponding allele frequencies of five major groups, Europeans (EUR), Africans (AFR), Americans (AMR), East Asians (EAS) and South Asians (SAS) (35). When conducting population analyses, the allele frequency is a key component, since it corresponds to the occurrence of a distinct genetic variant within a population (36). Allele frequencies, which are provided within the range of [0-1], constitute a reflection of genetic diversity; monitoring their changes allows the detection of shifts within the population (37).

As mentioned previously, COVID-19 exhibits variability across individuals, hinting at a trove of genetic factors which contribute to COVID-19 susceptibility and severity (38). As we wade through the third SARS-CoV-2 wave, the rapidly increasing volume of biomedical and genomic data calls for the implementation of modern techniques, for knowledge to be extracted and incorporated into novel therapeutic strategies. Natural language processing and other machine-learning techniques can make efficient use of the vast COVID-19 related literature, allowing the exploration of the complex architecture behind COVID-19 susceptibility and severity. Herein, we present a pipeline of semantic analysis of COVID-19 literature data for the mining of related SNPs, genes and disease ontologies. In the second phase of our analysis, we integrate population data from the 1000 Genomes Project. Our pipeline can serve as an example of an integrated approach in the research against COVID-19, towards estimating the “key” genomic target and providing beneficial knowledge in the personalization of medicine and the efficient assessment of populations at higher risk of infection and severe disease, on the basis of the genomic grammar and specifically SNPs they harbor.

## Methods

### Dataset collection and filtering

Using NCBI’s Entrez programming utilities, scientific literature in MEDLINE format was collected from Pubmed (39), limiting the search to the term “COVID-19” and publication dates post 2020. The MEDLINE files were collected in text form for subsequent filtering and feeding into the semantic analysis pipeline, which is summarized in Figure 1. A similar approach for the analysis of scientific publications has been described elsewhere (40). The filtering step of the pre-analysis included the removal of articles which were duplicates and unrelated to the subject.

**Figure 1.**
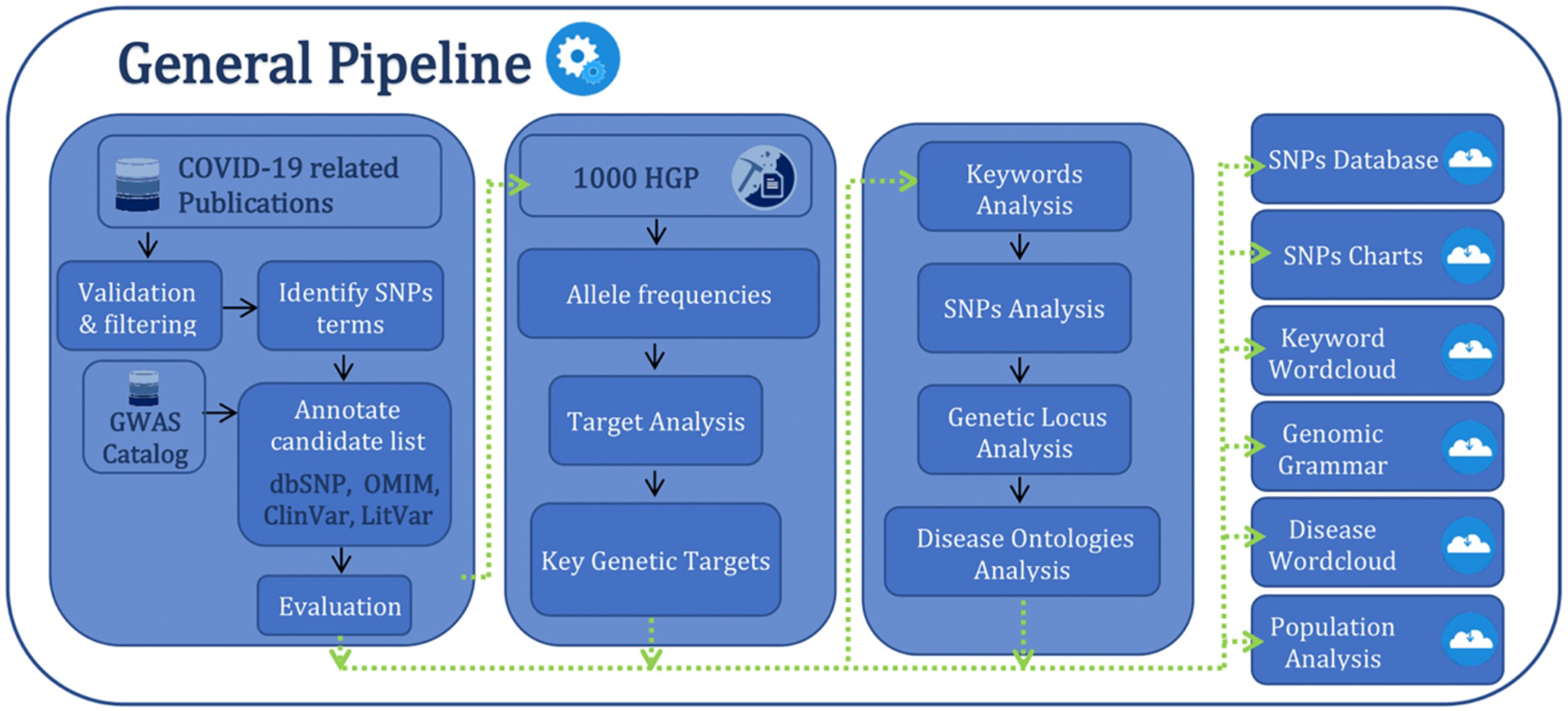
Summarized workflow of the study’s analysis.

### COVID-19 related SNPs

A search query was built with regular expressions in order to identify the candidate SNPs using the extracted dataset of the related articles with COVID-19. The extracted SNPs within the dataset of scientific articles were stored in a structured database for further analysis. Additionally, each article MEDLINE file was mined for supplementary information, such as MeSH/MEDLINE key terms, ontologies studied for their role in COVID-19 and mutations/polymorphisms. The candidate list was enriched with SNP data and meta-data from the GWAS Catalogue (41) and lastly, duplicates were removed from the candidate list.

### Data mining and semantics analysis

The candidate list of COVID-19 related SNPs was annotated through the use of publicly available databases of genetic variation, including dbSNP, ClinVar, LitVar and OMIM (42-45). The construction of the final SNPs dataset was carefully monitored; the annotated SNPs were evaluated, in order to remove those which were present as entries in the aforementioned databases and in the mined literature, but were actually reported as having no effect regarding COVID-19. Finally, the SNPs were subjected to semantics analysis to extract the desired knowledge, such as key terms, genomic grammar and disease ontologies (40, 46, 47).

### Population analysis

Five population groups were studied, focusing on Europeans, Africans, Americans, East Asians and South Asians. Sample sizes and origins of the individual population samples are given in the International Genome Sample Resource (IGSR) [78], which has been developed under the 1000 Human Genomes Project (1000 HGP) [79]. The elaboration of the present study has been performed using human genomes which were contained in the phase three collection of the IGSR on reference assembly GRCh38 (48, 49). Since the 1000 Genomes Project has created call sets of sequence variants for each of the different genomes sequenced, the downloaded data were multi-individual VCFs (50) per chromosome, with genotypes listed for each sample (49). Histograms regarding each population were generated with suitable packages of the Python programming language. Statistical analysis of the results was carried out with the use of the R Biocircos package, which enables the visualization of genomic-related data and is based on the Javascript library developed by Cui et al (51).

## Results

### COVID-19 related key terms and SNPs

The collection of COVID-19 related biomedical literature enables the identification of keywords as they appear within the MEDLINE files. Through querying of the Pubmed database, 147.396 non-duplicate scientific articles corresponding to the search term “COVID-19” were collected in a final dataset and were subsequently mined for related keywords. A total of 98.497 keywords were assembled, out of which 2.677 were identified as most frequent, providing a first estimation of relation to COVID-19. The most frequently appearing keywords are visualized as a word cloud in Figure 2. The word cloud visualization technique enables the presentation of the results, where the size of the words – in this case the keywords – indicates their frequency within the dataset. SNPs were found to be contain within 147 of the collected articles. Following their extraction, their enrichment through GWAS Catalogue and their annotation, a total of 526 SNPs were collected, to be further subjected to evaluation. Out of them, 339 SNPs related to COVID-19 were identified and collected.

**Figure 2.**
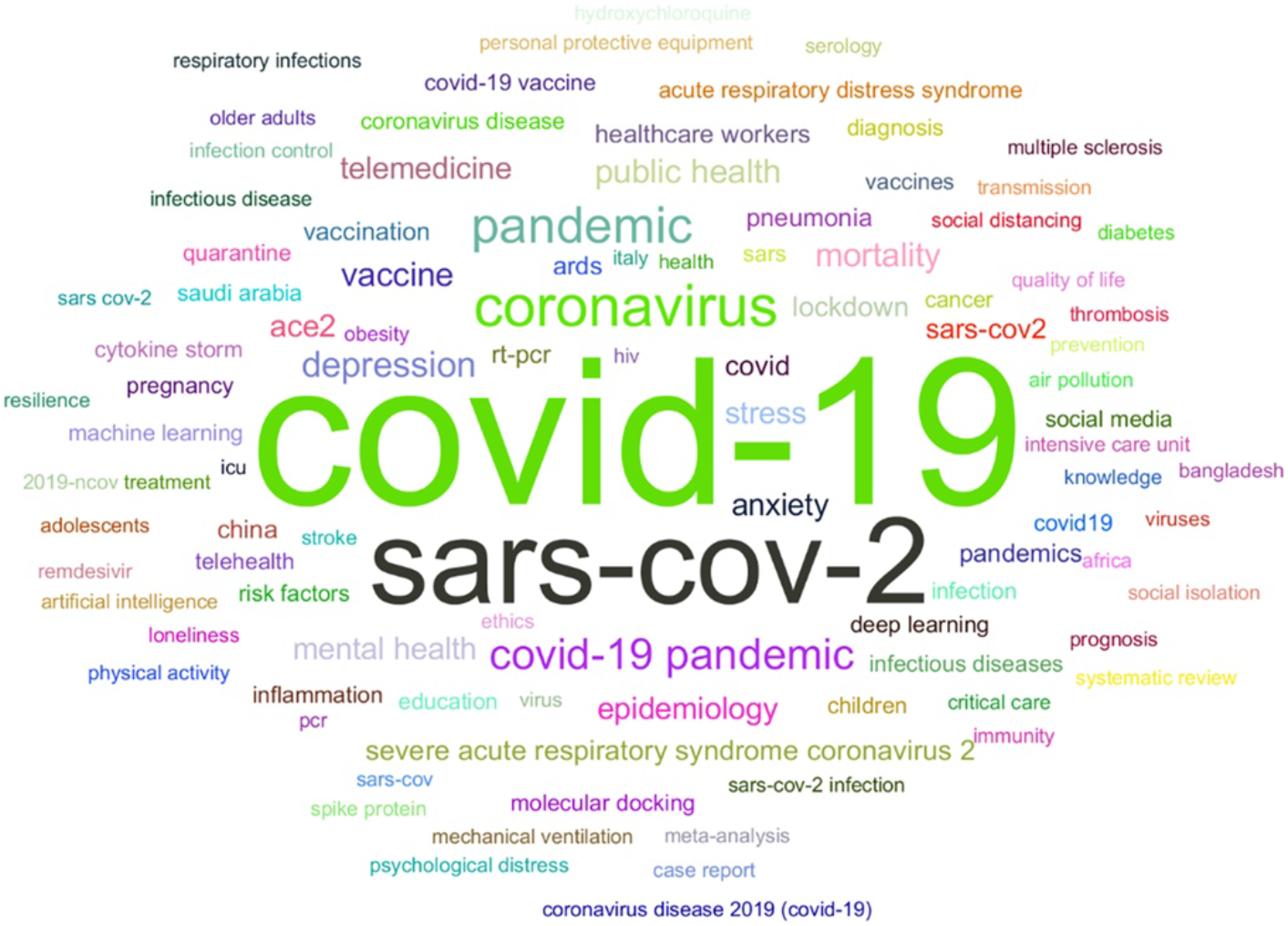
Word cloud presentation of COVID-19 related keywords, according to the input dataset of literature articles. The size of the words in the cloud mirrors their frequency within the dataset.

### Genomic grammar

After semantics analysis of the annotated and evaluated COVID-19-related SNPs, the genomic grammar of the disease was constructed. The genomic grammar, as mentioned earlier, constitutes the genomic map of COVID-19 variation, encompassing the genes and non-coding locations which harbor the related SNPs. The results concerning COVID-19’s genomic grammar are visualized in word-cloud form in Figure 3. Since the size corresponds to the frequency of the genomic “word”, a quick survey of Figure 3 enables the identification of some of the prominent genetic players related to COVID-19. For example, *ACE2* is easily identified as a central gene. In the renin-angiotensin system (RAS), which is important for blood pressure levels and by extension for the proper function of multiple organs, angiotensin I (Ang-II) and angiotensin II (Ang-II) constitute important biomolecules (52). The angiotensin-converting enzyme (ACE) converts Ang-I to Ang-II, which can bind to angiotensin type I receptor (AT1R) and angiotensin type II receptor (AT2R) (53). Interaction between Ang-II and AT1R triggers processes such as vasoconstriction, fibrosis and inflammation, while interaction with AT2R counteracts the AT1R-mediated effects (54). ACE2 is a homologue of ACE and can convert Ang-I and Ang-II to angiotensin-(1-7) (55). Angiotensin-(1-7) acts on the G protein-coupled receptor MAS to carry out processes such as anti-fibrosis, anti-inflammation, generally having an effect opposite to that of Ang-II/AT1R binding (56). As previously mentioned, ACE2 participates in the entry of SARS-CoV-2 into the host cells through interaction with the viral surface protein S (57). Significant ACE2 increase has been documented in patients with severe COVID-19 (58), and since this receptor is highly expressed in various organs and tissues and is a major component of inflammation, its association with multiple-organ failure syndromes in COVID-19 remains under intense study (59, 60). *ACE2* polymorphisms and their effect on COVID-19 severity, susceptibility and progression are gradually being explored as a tool to monitor the disease outcome on an individual patient level (61-63).

**Figure 3.**
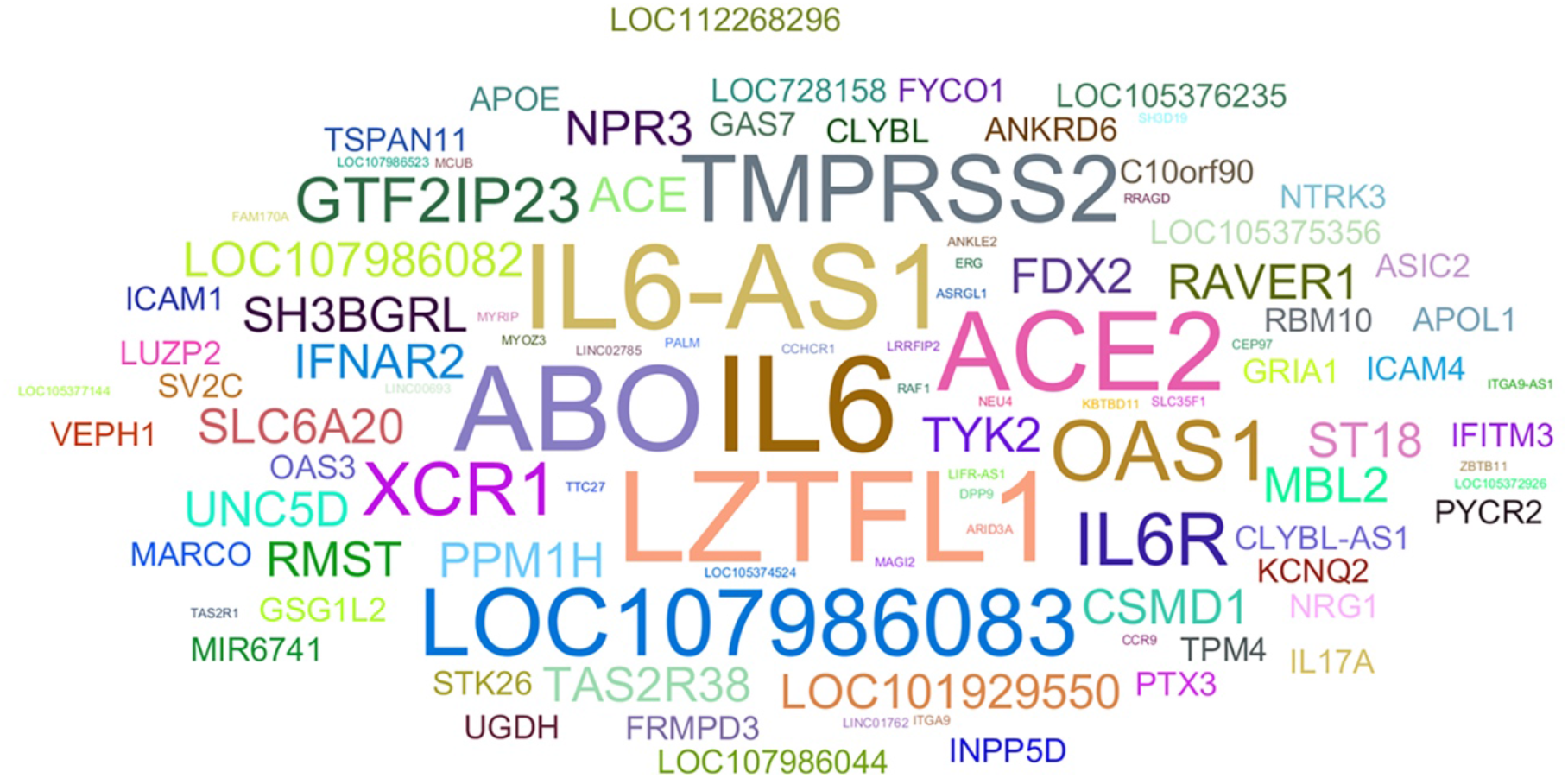
Word-cloud presentation of COVID-19 related genes and other non-coding regions. The size of the words in the cloud corresponds to their frequency within the respective dataset. Prominent genes and non-coding regions such as ACE2, ABO, IL-6, TMPRSS2, ZTFL1 and LOC107986083 can be identified at a first glance.

Interleukin-6 (IL-6) is a cytokine with multifaceted involvement in the initiation of immune response and inflammatory processes (64). Viral infection triggers its secretion by immune cells such as macrophages, B and T cells, as well other cell types such as endothelial cells and fibroblasts (65). IL-6 is a critical component of the cytokine storm phenomenon in cases of severe COVID-19 and high levels of IL-6, among other cytokines, are one of the hallmarks of severe COVID-19 (66, 67). Polymorphisms in the promoter and regulatory regions of the *IL-6* gene modulate the protein’s expression, helping to account for the differences in immune response documented in various ethnic groups against a variety of pathogens, including SARS-CoV-2 (68-70). IL6-AS1 (IL6 Antisense RNA 1) is a long non-coding RNA and has been found to be upregulated in chronic obstructive pulmonary disease, promoting the expression of IL-6 (71). An *IL-6* variant haplotype common in Asian populations was found to be protective against severe COVID-19, associated with lower IL-6 and IL-6-AS1 levels through a disturbance of a binding locus at the IL-6-AS1 enhancer elements (69).

Transmembrane serine protease 2 (TMPRSS2) is an essential host factor for SARS-CoV-2 pathogenicity and an important player in the viral entry into host cells, priming the viral S glycoprotein for viral fusion (72).

*TMPRSS2* SNPs have been the object of study for the establishment of disease outcome biomarkers in pathologies such as cancer and severe viral infections such as H1N1 infection (73, 74). A computational analysis aimed at explaining susceptibility differences among populations identified a number of SNPs with predicted effect on protein function (75), while a study in print elected *TMPRSS2* genetic variants as candidate COVID-19 modulators after examining single nucleotide polymorphisms in various ethnic populations (76). Lastly, a common *TMPRSS2* non-synonymous variant, rs12329760, was found to be protective against severe COVID-19 through impacting the enzyme’s catalytic ability and thus its role in the viral entry (77).

Variations in the *ABO* gene in chromosome 9 are the basis for the establishment of the conventional ABO blood group (78). The *ABO* locus encodes three alleles, with alleles A and B producing α-1,3-N-acetylgalactosamine transferase, α-1,3-galactosyl transferase B respectively, while allele O exhibits a deletion-caused frameshift and lacks both of the aforementioned enzymatic activities (79). Increased levels of ABO protein in plasma appear to be associated with risk of severe COVID-19 (80). The same study linked COVID-19 risk and severity with the OAS1 gene, which is activated by interferon and participates in the cellular innate antiviral response (81). Although the exact mechanism underlying the effect of the ABO blood group on COVID-19 susceptibility and severity remains under investigation, a recent study reported association of blood groups A and B with increased risk of SARS-CoV-2 infection (82), while another study found B-allele frequencies to be correlated with COVID-19 mortality (83).

The *LZTFL1* gene codes for a leucine zipper protein, which associates with E-cadherin and participates in the circulation of a variety of signaling molecules (84, 85). The gene is expressed in pulmonary epithelial cells, among others, and has been recently identified as a target for a probable causative variant related to COVID-19 risk (86, 87). Rs17714054A, the risk allele of the SNP, was found to target *LZTFL1*’s enhancer region, leading to the gene’s upregulation (86). According to NCBI data, LOC107986083 is an uncharacterized non-coding RNA located in chromosome 3 and has been found to be broadly expressed in the testis and thyroid, among other tissues. Positionally, it is associated with the LZTFL1 gene.

### Disease ontologies

The semantics analysis of the SNPs, related genes and non-coding regions, enables the subsequent extraction of information regarding disease profiling related to COVID-19. The diseases identified through our analysis are summarized in world-cloud form in Figure 4, where the size of the words, or disease terms in our case, mirrors the frequency of the term. A number of prevalent words, such as neoplasms, chronic hepatitis C or obesity, can thus be easily identified with a first study of the visualized results. Neoplasms are abnormal and excessive tissue growths which, when malignant, are known as cancers (88). Patients with hematologic malignancies were found to be in a significant risk of COVID-19 related death (89). In a preliminary, exploratory analysis, essential thrombocythemia, a type of myeloproliferative neoplasm, was found to be associated with greater risk of venous thromboembolism in COVID-19 patients (90). Similarly, myeloproliferative neoplasm patients and especially those suffering from myelofibrosis were in high risk of mortality during COVID-19 infection (91). Paraoxonase-1 (PON1) is a lactonase which degrades lipid peroxides in low-density lipoproteins (LDL) and high-density lipoproteins (HDL) (92). The PON1-L55M polymorphism at the *PON1* genetic locus has been identified as a potential risk factor for breast cancer (93). Furthermore, *PON1* genetic variants are under investigation for involvement in predisposition to obesity-associated fatty liver disease (94). A recent study investigating a potential association between *PON1* polymorphisms and COVID-19 pointed at a possible positive correlation between M55 and prevalence and mortality of COVID-19, although these findings remain to be validated in larger-scale studies (95).

**Figure 4.**
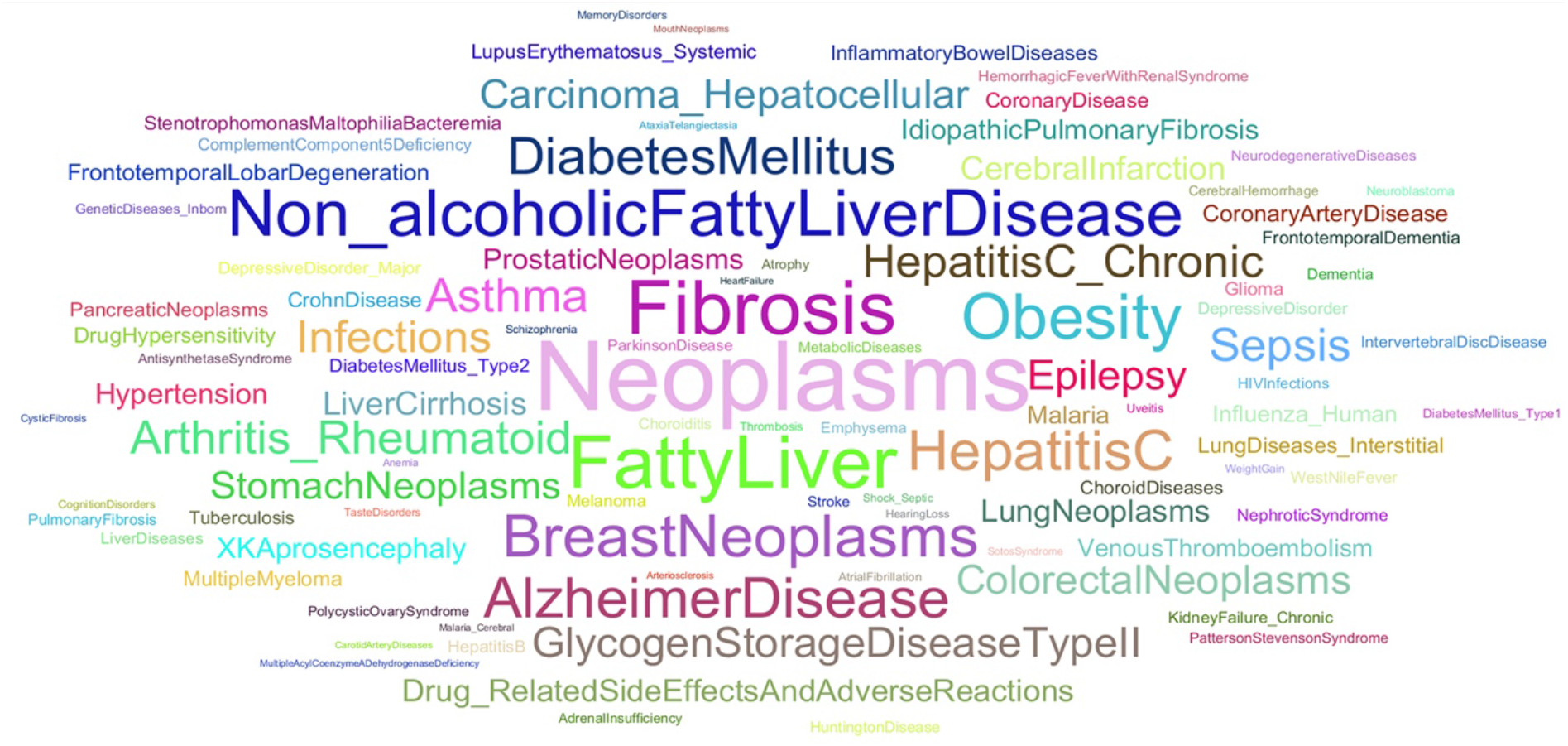
Word-cloud presentation of disease ontologies related to COVID-19. Word size within the cloud reflects the frequency of the word, allowing the quick identification of strongly related diseases, such as neoplasms, non-alcoholic fatty liver disease, fibrosis and obesity.

Increased left ventricular strain as a fibrosis marker was found to predict moderate and severe COVID-19 progression with accuracy (96). Myocardial fibrogenesis is induced by transforming growth factor β1 (TGF-β1) (97, 98). An increase in TGF-β1 expression possibly leads to susceptibility to SARS-CoV-2 infection as a result of increased expression of NRP-1 neuropilin-1 (NRP-1), a co-receptor for TGF-β1, in the lungs (99). Lastly, lung inflammation and pulmonary fibrosis can be potentially promoted by SARS-CoV-2 complications, such as Acute Respiratory Distress Syndrome (ARDS) (100).

Non-alcoholic fatty liver disease (NAFLD) is a condition marked by fat accumulation in the parenchymal region of the liver without ties to hepatic fat accumulation factors, such as alcohol abuse or hepatitis B infection (101). NAFLD patients are susceptible to COVID-19 due to an existing layer of risk factors, such as type 2 diabetes, which promotes susceptibility to infection (102), obesity, which is linked to respiratory complications (103), and cardiovascular disease (104). A meta-analysis study between NAFLD and non-NAFLD patients reported an increased risk of severe COVID-19 infection and ICU admission (105). The added presence of obesity in NAFLD patients appears to increase the severity of COVID-19 (106). In a pooled study of COVID-19 and NAFLD data, the presence of NAFLD was associated with an increased risk of severe COVID-19 (107). Obesity has been associated with various inflammatory mediators such as IL-6 (108, 109), which in our study was found to be part of COVID-19’s genomic grammar. Subsets of immune cells in the white adipose tissue lead to a surge in inflammation-promoting cytokines like tumor necrosis factor α (TNFα) and IL-6 (108). TNFα and IL-6 are players in the signaling of the initial phase of cytokine-storm, a phenomenon prevalent in severe COVID-19 (110, 111). The hyperinflammatory state observed in obese individuals may also lead to coagulopathies (112), the hallmark of which are shifts in the levels of D-dimer, a fibrin degradation product (FDP) (113). Correlation has been shown between D-dimer and COVID-19 severity (114).

Chronic hepatitis C is caused by the hepatitis C virus (HCV) following acute infection, with potential complications such as liver damage, cirrhosis and cancer (115). Genetic variation at the Interferon lambda 4 (IFNL4) genetic locus has been studied with the aim to identify viral clearance predictors (116). One polymorphism, rs12979860, has been associated with clearance of hepatitis C virus and other RNA viruses which target the upper respiratory system (117, 118). Additionally, association has been evidenced between this polymorphism and the response to type I IFN treatment efforts in patients with chronic hepatitis C. The T allele of the aforementioned polymorphism was found to be overexpressed in COVID-19 patients, highlight its potential as a risk factor for COVID-19 (119).

Apolipoprotein E (APOE) e4 genotype, which has been linked to high risk for Alzheimer’s disease, was described as a potential predictor of severe COVID-19 infection (120). In addition, a study by Taylor *et al*. identified four severe COVID-19 risk-associated genes which had been previously linked to increased risk of developing Alzheimer’s, further hinting at possible interplay between the two diseases (121).

### Population analysis

Directional change and reversal in allele frequencies has been shown in the 339 COVID-19 related SNPs between the individuals of the major five clusters. The histogram analysis of COVID-19 related SNPs using the 1000 Genome Project dataset shows similar distribution in the five major groups (Figure 5). Although different allele frequencies have been identified in the studied SNPs, some groups appear to have similar distributions with different numbers such as the Africans and East Asians example or the Europeans and Americans example (Figure 5). The five studied groups have accumulated different SNPs totals at the sensitive two extremities of the allele frequencies including the cluster of the “low allele frequencies” (0.1 ≥ SNP allele frequency) and the cluster of the “high allele frequencies” (0.9 ≤ SNP allele frequency) (Figure 2) (122-124). Our findings are in agreement with the expected observations (125).

**Figure 5.**
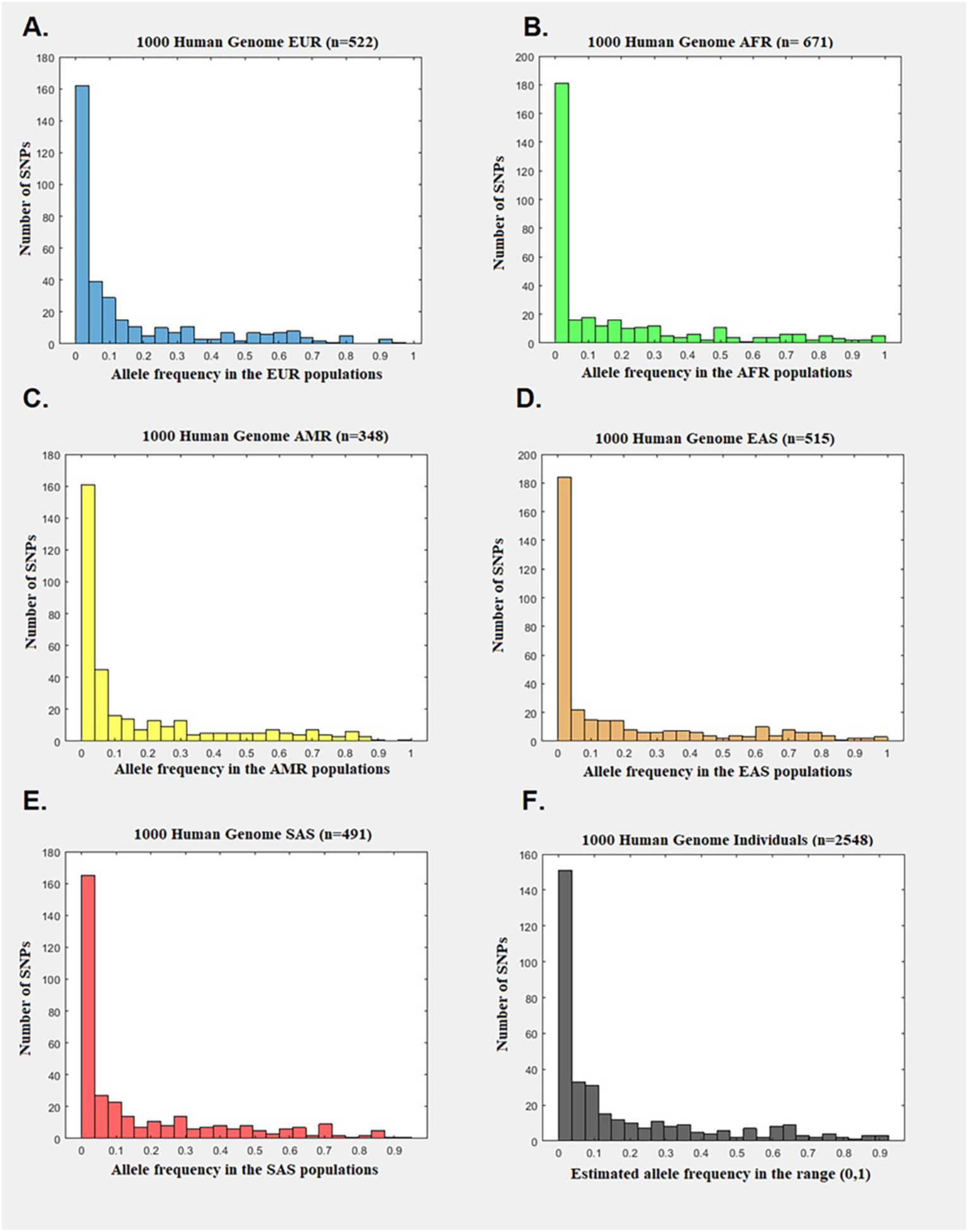
Histogram analysis of the Histogram of the allele frequencies of the COVID-19 related SNPs as extracted from the 1000 Human Genomes project. (A) Histogram of the SNPs allele frequencies for the separated group of the Europeans. (B) Histogram of the SNPs allele frequencies for the separated group of the Africans. (C) Histogram of the SNPs allele frequencies for the separated group of the Americans. (D) Histogram of the SNPs allele frequencies for the separated group of the East Asians. (E) Histogram of the SNPs allele frequencies for the separated group of the South Asians. (F) Histogram of the SNPs allele frequencies for total number of the studied individuals.

Different totals and ids of SNPs have been accumulated in the low and high clusters between the population groups (Figure 5). The American group has the largest sample of SNPs with low allele frequencies followed by Europeans, South Asians, East Asians, and Africans (Figure 5 C, A, E, F, B). On the other hand, the African group has the largest sample of SNPs with high allele frequencies followed by the East Asian group (Figure 5 B, D). The South Asian, American and European groups show significant fewer totals in SNPs with high allele frequencies (Figure 5 E, C, A). Although some population groups are shown some similarities in the ids of the identified SNPs in the low and high clusters, the overall distribution of the COVID-19 related SNPs and their genetic locus per chromosome in the five studied population groups shows a significant differentiation. A general conclusion to be drawn from the results is that the genomic grammar of Africans and East Asians contains more COVID-19 related SNPs in the two sensitive clusters (low and high) than the other groups (Figures 5,6).

**Figure 6.**
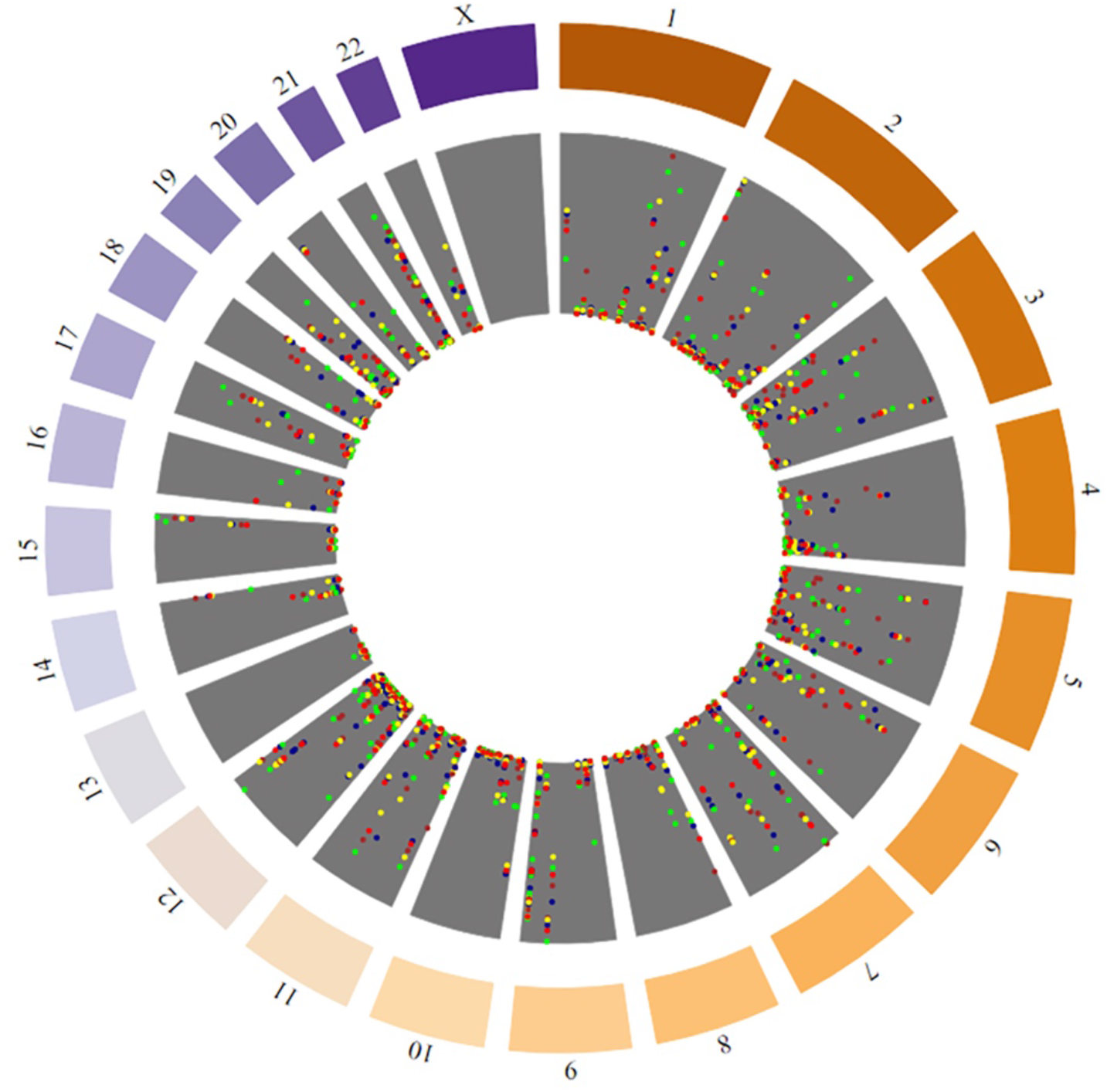
Circos-like visualizations of the genomic data of the five major population groups in the identified COVID-19 related SNPs with low allele frequencies (Blue dots = Europeans, green dots = Africans, yellow dots = Americans, brown dots = East Asians and red dots = South Asians).

COVID-19 related SNPs at the high allele frequency cluster were identified for each population and trends in their distribution were studied. Table 1 summarizes results regarding the high-frequency cluster. The largest number of COVID-19 related SNPs with high allele frequency can be observed within the African population, while the smallest number can be observed within the American population. Europeans and East Asians appear to share a similar sum of COVID-19 related SNPs with high allele frequency, although there is distinction between the specific high-frequency allele SNPs which they harbor.

**Table 1.** Distribution of the COVID-19 related SNPs of the high allele frequencies across the five studied population groups.

| SNP name | EUR [A] | AFR [B] | AMR [C] | EAS [D] | SAS [E] |
| --- | --- | --- | --- | --- | --- |
| **rs6775748** | ✓ | ✓ |  | ✓ | ✓ |
| **rs6054661** | ✓ |  |  |  |  |
| **rs11127334** | ✓ | ✓ | ✓ | ✓ |  |
| **rs2795384** | ✓ | ✓ |  |  | ✓ |
| **rs4766676** |  | ✓ |  |  |  |
| **rs1800795** |  | ✓ |  | ✓ |  |
| **rs1800797** |  | ✓ |  | ✓ |  |
| **rs4702** |  | ✓ |  |  |  |
| **rs1126579** |  | ✓ |  |  |  |
| **rs180040** |  | ✓ |  |  |  |
| **rs6127** |  |  |  | ✓ |  |

Population-specific percentages of SNPs of the high and low allele frequency class against the total number of COVID-19 related SNPs are presented in Table 2. It can be observed that low-frequency SNP alleles exhibit a high percentage - more than 60% across all populations - while high-frequency range between 0,5 and 2,7%. These results could be the basis for designing a common treatment for endometriosis with significant discrepancies depending on the population group (126, 127).

**Table 2.** Percentage of COVID-19 related SNPs with high and low allele frequencies within the five studied populations.

| **Population** | ***High %*** | ***Low %*** |
| --- | --- | --- |
| EUR [A] | 1,17 | 67,25 |
| AFR [B] | 2,65 | 63,12 |
| AMR [C] | 0,29 | 64,30 |
| EAS [D] | 1,47 | 64,60 |
| SAS [E] | 0,58 | 62,24 |

GWAS Catalogue reports our “high”-SNP rs6775748 as an intergenic variant, which was identified in a study investigating genetic and nongenetic COVID-19 associations (128). Genomic regions closest to the intergenic variant include the *SI* gene, which codes for the sucrase-isomaltase protein, and LINC01324, a long intergenic non-protein coding RNA (ncRNA) 1324 (129-131). The same study which reported rs6776748 also reported rs6054661, another intergenic variant mapped between the *BMP2* gene, which codes for bone morphogenetic protein 2, and LINC01428, a long intergenic ncRNA (128). Rs11127334 is an intron variant which is mapped to *MYT1L*, the gene which codes for myelin transcription factor 1 like protein, and the intergenic lncRNA LINC01250 (128, 132). Rs2795384 constitutes an intergenic variant, and its mapped genomic regions include *ARL2BPP7*, the ADP ribosylation factor like GTPase 2 binding protein pseudogene, and *MTND3P4*, MT-ND3 pseudogene 4 (128, 133). Rs4766676 is mapped in the OAS1 gene, more specifically as an intron variant. Intronic variants can influence the process of alternative splicing through interference with the recognition of the splice site, potentially leading to production of malfunctioning protein products (134). As mentioned, 2′-5′-oligoadenylate synthetase 1 (OAS1) is a key player against viral infections, as this interferon-activated enzyme degrades viral RNA in partnership with RNase L (135). According to GWAS Catalogue, rs1800795 is an intron variant of the *IL6* gene, its antisense RNA 1 (*IL6-AS1*), and *STEAP1B*. The latter may encode two different transcripts, STEAP1B2, which is overexpressed in prostate cancer cells, and STEAP1B1 (136). *IL6, IL6-AS1* and *STEAP1B* are also mapped to the rs1800797 polymorphism, a non-coding transcript exon variant. This variant has been linked to two immune-related pathologies, asthma and systemic lupus erythematosus (137, 138). Rs4702 is a variant located in the 3 prime untranslated region (UTR) of the *FURIN* gene, which codes for FURIN, a pro-protease convertase bound to host membranes (139). This SNP has been shown to influence alveolar and neuron infection by SARS-CoV-2 *in vitro* (140). The SARS-CoV-2 spike harbors a FURIN cleavage site, which promotes entry into lung cells and is absent from SARS-CoV (141, 142). Rs1126579 is an SNP identified at the *CXCR2* gene, leading to a 3 prime UTR variant. The gene codes for C-X-C motif chemokine receptor 2 (CXCR2), a key stimulator of immune cell migration, which binds to interleukin 8 (IL8) and chemokine ligand 1 (143). Another “high” SNP, rs180040, is mapped to the *CYP1B1* gene, which encodes an enzyme of the cytochrome P450 family of monooxygenases that catalyze reactions of lipid synthesis and drug metabolism (144). Drug clearance is an important element in COVID-19 patient treatment; the state of hyperinflammation which is often observed in COVID-19 can potentially alter the function of cytochrome P450 enzymes in critical organs, thus affecting drug clearance and the course of therapeutic regiments (145). Lastly, rs6127 is mapped to the *SELP* gene, which codes for selectin P, a cell adhesion molecule (146). In a recent whole exome sequencing study, the polymorphism was found to be associated with thrombosis and COVID-19 severity in male patients (147). Overall, the COVID-19 related SNPs which fall into the cluster of high allele frequency are located in varying genomic regions, from introns and exons to intergenic regions. These results provide insight into key genetic targets within the studied population groups, with the potential to inform and guide policies of management and treatment according to the population-specific COVID-19 related SNPs and their corresponding clusters of allele frequencies.

## Discussion

Human SNPs and their influence on enhanced resistance or susceptibility to viral disease have been the subject of intensive research, applied to pathogens of global concern, such as influenza or HIV (148-150). Similarly, human genomic variants related to COVID-19 severity and sensitivity to infection are being evaluated as tools to guide and adjust therapeutic strategies, such as the choice of administered drugs (151). The wide range of pathologies that already exist within the global population inevitably lead to the formation of a complex web, with layers of potential comorbidities between them and COVID-19. As we move into the age of “omics” and the COVID-19 related data becomes vast and heterogeneous, the modern framework of computational systems lends itself to researchers as a powerful tool. The natural-language processing pipeline proposed herein enables the effective search for potential connections between COVID-19, genes and other diseases, using the trove of characterized SNPs as our guiding light. With a combination of semantic analysis and machine learning we have drawn COVID-19’s “genomic grammar”, i.e the associated genomic regions which house the SNPs. Furthermore, we have examined disease profiling ontologies in connection with COVID-19, such as neoplasms, chronic hepatitis C and Alzheimer’s. This firstly allows the identification of risk groups and secondly, may inform efforts towards personalized medicine, where the patient’s genomic makeup determines the therapeutic approach. Lastly, we sought to expand our results to a populational scale, encompassing data from the 1000 Genomes Project, to gain insight into key genetic targets for potential exploration in studied population groups. A major portion of the scientific community’s effort is dedicated to studying SARS-CoV-2’s genome and its coding products in the search of effective ways to control and face the ongoing pandemic. Simultaneously, the genetic background of the human host – and its variations - constitute a source of invaluable knowledge, which can complete the picture and inform the search for effective, safe and targeted COVID-19 therapeutics.

## Acknowledgments

Not applicable.

